# Novel inhibitors designing against the potential drug target of *Plasmodium falciparum* M17 Leucyl Aminopeptidase – PfM17LAP

**DOI:** 10.1101/2021.12.14.472562

**Authors:** Govinda Rao Dabburu, Manish Kumar, Naidu Subbarao

**Affiliations:** Department of Biophysics, University of Delhi South Campus, New Delhi-110021, India; School of Computational and Integrative Sciences, Jawaharlal Nehru University, New Delhi 110067, India

**Keywords:** Malaria, M17 Leucyl Aminopeptidase, ADMET, X-SCORE

## Abstract

Malaria is one of the major disease of concern worldwide especially in the African regions. According to the recent WHO reports, African regions share 95% of the total deaths worldwide that occurs due to malaria. *Plasmodium falciparum* M17 Leucyl Aminopeptidase (PfM17LAP) plays an important role in the regulation of amino acids release and for the survival of the parasite. We performed molecular docking and simulation studies to find the potential inhibitors against PfM17LAP using ChEMBL antimalarial library. Molecular docking studies and post-docking analysis revealed that molecules CHEMBL369831 and CHEMBL176888 showed better binding than the reference molecule BESTATIN. LibDock and X-SCORES of molecules BES, CHEMBL369831 and CHEMBL176888 are 130.071, 230.38, 223.56 and -8.75 Kcal/mol, -10.90 Kcal/mol, -11.05 Kcal/mol respectively. ADMET profiling of the top ten ranked molecules was done by using the Discovery Studio. Molecular dynamic studies revealed that the complex PfM17LAP-CHEMBL369831 is stable throughout the simulation. Finally, we have reported novel inhibitors which possess more binding affinity towards PfM17LAP.

## Introduction

Malaria is one of the most lethal diseases transmitted by the *Plasmodium* spp through bite of female Anopheles mosquito. *Plasmodium falciparum* is one of the most lethal apicomplexan parasites among the *Plasmodium* species. Deaths due to malaria are more in countries with poor sanitization and in African countries. Children under age of 5 years are more prone to the malarial infection in African regions, along with that African regions shares 95% of the malarial cases and 96% of the overall malarial deaths throughout worldwide. According to the WHO statistics, there are 229 million malaria cases and 409000 deaths reported worldwide in 2019. There are 241 million malaria cases and 627000 deaths reported worldwide in 2020. Deaths due to malaria increased in 69000 number as compared to the last year 2019. In October 2021, WHO approved the use of the malarial vaccine RTS,S/AS01 for children in nations with high transmission rate of *P. falciparum*. To achieve the success in the malarial treatment Artemisinin is being used either as a single drug or in combination with other drugs (ACT), use of Artemisinin in combination with other drugs helps to overcome the resistance against the other drugs. Other advantages of ACT are high efficacy, fast action and likelihood of resistance development.

ACT therapies are showing better results in the terms of efficacy and its control over the malaria is good. But in the recent years ACT therapy is also showing resistance in some areas of the Western Cambodia (Leang et al. 2015) and failure to the ACT therapies are rapidly increasing (Mok et al. 2011). Artemisinin resistance is emerging and spreading independently in the mainland Southeast Asia, Pursat, Western boarder of Thailand, Southern Myanmar and Vietnam (White et al. 2014)(Kyaw et al. 2013). A single nucleotide polymorphism in Kelch 13 gene of the *Plasmodium* parasite is reported as the causes of resistance related to Artemisinin (Noreen et al. 2021).

Malaria parasite utilizes proteases in the many stages of its life cycle. Aminopeptidase enzymes remove N-terminal amino acids from short peptides with high specificity. Two Aminopeptidases namely *P. falciparum* M1 Aminopeptidases (PfM1AAP) and *P. falciparum* M17 Leucyl Aminopeptidase (PfM17LAP) have important role in the growth and development of parasites inside the red blood cell by regulating release of amino acids that are required for the growth of parasite (Ruggeri et al. 2015). These two enzymes are present in the cytosol of parasite and are responsible for the survival of parasite. They also provide nutrients required for the growth and development of parasite. Gene targeting and in vivo experimental results have shown that, inactivation of PfM17LAP causes inhibition of growth in some parasites and in some parasites death also (Stack et al. 2007)(McGowan et al. 2009).

Expression data shows that PfM17LAP expression in all stages of Plasmodium life cycle namely intra-erythrocytic, merozoite, sporozoite, early ring, early schizont, late schizont, late trophozoite, oocyst and ring stages (Lasonder et al. 2016)(Otto et al. 2010)(Zanghí et al. 2018).

Hence we used Insilico tools to develop novel inhibitors of PfM17LAP with the aim to stop growth of parasite. PfM17LAP is 528 amino acid structure with two metals in the active site. The metal of the first site Mg^2+^ is readily exchangeable with other metal ions Zn^2+^, Co^2+^ and Mn^+2^ and acts as a regulatory site. But the second site with Zn^2+^ is tightly bound to the active site and known as catalytic site. Metal replacement studies found that PfM17LAP retains activity when the tight binding catalytic site is occupied with the metal ion, but removal of the both metal ions leads to the irreversible inactivity (McGowan et al. 2010).

The structure of the PfM17LAP shows that only catalytic site 2 occupied by the Zn^2+^ and no electron density was visible for the site occupied by Mg^2+^. Zn^+2^ coordinated by the Lys374, Asp379, Asp399 and Glu461. Along with the metal ions it also have inhibitors BESTATIN (BES) and Co4. BES is natural inhibitor of the leucine aminopeptidases. It is obtained from the culture filtrates of the *Streptomyces olivoreticuli* (Wilkes and Prescott 1985). It has antibiotic and antimalarial activities as well (Wilkes and Prescott 1985). Inhibitory activity (Ki) for the BES and Co4 are 25 nM and 13 nM respectively (McGowan et al. 2010).

*Plasmodium* parasite uses Plasmepsins for intraerythrocytic hemoglobin degradation of human host and uses it as nutrient source (Kumar and Ghosh 2007). There are 10 different types of the Plasmepsins in the *Plasmodium* species expressed at different stages of the life cycle. Among the 10 types, PfPM I-IV are active in digestive vacuole and shares 50-70% identity; PfPM-V active in effector export and it shares 19-23% identity with other PfPM; PfPM-VI, VII & VIII are active during transmission stage and they shares 31-36% identity among each other; PfPM-IX and X shares 37% identity between each other (Nasamu et al. 2020). PfPM III is also known as Histidine Aspartic Protease (HAP), due to presence of Histidine in the active site. Expression data of the PfPM I-IV are mentioned in table 1. Studies have shown that removal of the any one of the PfPM from PfPM I-IV does not affect the parasitic activity but removal of multiple PfPMs show effective antiparasitic activity (Liu et al. 2005). Hence considering the importance of PfPM I-IV in parasitic activity, we selected PfPM I-IV for molecular docking studies. Expression rate of PfPM I-IV in the different stages of *Plasmodium* life cycle and their corresponding PDB IDs are mentioned in table 1.

**Table 1:** Expression data of Plasmepsins I-IV from TDR targets along with the corresponding PDB IDs from Protein Data Bank. Structures highlighted in bold color are used for docking in this study. HAP-Histidine Aspartic Protease.

| Plasmepsins | Expression data from TDR targets (80-100%) | PDB structures |
| --- | --- | --- |
| Plasmepsin I / PfPM-I | Intra-erythrocytic, merozoite, sporozoite, early ring, early trophozoite, late ring, late schizont and ring stages | 3QRV, **3QS1** |
| Plasmepsin II /<br>PfPM-II | Intra-erythrocytic, merozoite, early trophozoite, late ring and ring stages | 1J8J, 1LEE, 1LF2, 1LF3, 1LF4, 1M43, 1ME6, 1PFZ, 1SME, 1W6H, 1W6I, 1XDH, 1XE5, 1XE6, **2BJU**, 2IGX, 2IGY, 2R9B, 3F9Q, 4CKO, 4CKU, 4Y6M, 4YA8, 4Z22, 5BWY, 5YIA, 5YIB, 5YIC, 5YID, 5YIE |
| Plasmepsin III / HAP / PfPM-III | Intra-erythrocytic, gametocyte, merozoite, sporozoite, early ring, early schizont, early trophozoite, late ring, late schizont, late trophozoite and ring stages | 3FNS, 3FNT, 3FNU, **3QVC**, 3QVI, 6KUB, 6KUC, 6KUD |
| Plasmepsin IV / PfPM-IV | Intra-erythrocytic, merozoite, sporozoite, early ring, early schizont, early trophozoite, late ring, late schizont, late trophozoite and ring stages | **1LS5**, 1MIQ, 1QS8, 2ANL, 5I70, 5JOD |

To design novel inhibitors against PfM17LAP we performed molecular docking studies. We selected CHEMBL antimalarial library for screening, which have bioactivity information for every molecule. We selected top ten molecules with high docking score for post docking analysis and for Molecular Dynamic (MD) simulations.

For the selected the targets PfPM I-IV, to design the potential antimalarial drugs. We performed molecular docking and post-docking analysis of the top ten hits obtained from the high throughput screening of the ChEMBL database against the PfM17LAP, to propose the dual inhibitory activity of the top ten hits.

## Material and Methods

### Protein preparation and chemical library preparation

PfM17LAP crystal structure is taken from the Protein Data Bank (PDB ID - 3KR4). Resolution of PDB structure is 2A^0^. Active site residues are defined based on the reference ligand BES. PDBSum analysis showed that residues ASP379, LYS386, ASP399, ASP459, GLU461 and GLY489 makes hydrogen bonds with BES.

Structures of PM-I (PDB ID - 3QS1), PM-II (PDB ID - 2BJU), PM-III (PDB ID – 3QVC) and PM-IV (PDB ID - 1LS5) were also retrieved from PDB database.

Proteins are prepared and energy minimized by using the protein preparation and minimization wizards of the Discovery studio. Loops are minimized by using CHARMm forcefield. Prepared proteins are minimized by using the Smart Minimizer to get the lowest possible energy conformation of the protein.

ChEMBL antimalaria database was used for the screening purpose. ChEMBL antimalarial database had 282295 bioactive compounds. ChEMBL database is a collection bioactive compounds from different assays reported in literature and from other sources. Compounds are prepared by using Discovery studio. A maximum of 10 tautomers were generated for each compound. Isomers were generated and bad valancies were fixed during the library preparation. 3D co-ordinates were generated for each compound. A total of 281896 compounds generated after library preparation.

### Molecular Docking Studies

LibDock module of Discovery studio was used for molecular docking; LibDock is flexible docking method that uses high throughput molecular docking algorithm developed by Dille and Merz (Diller and Merz 2001). A total of 281896 compounds were taken for molecular docking studies. Binding site sphere was defined with radius 87.7078, 73.2827, 30.8208 and 11.1426. FAST ligand conformation method was used for generating ligand conformations. 255 conformations are generated for every compound. LibDock uses polar and apolar hotspots to generate the protein site features. Top 100 scored conformations of each compound are saved after docking process.

### Post-docking analysis

Top ten LibDock scored compounds were selected for the post-docking analysis. Protein-ligand interactions were calculated and visualized by using Ligplot+ (Laskowski and Swindells 2011). Ligplot generates interaction diagrams of hydrogen and non-bonded interactions between the protein-ligand complexes along with labelling of distance between the bonds. Binding energies between receptor and ligand were calculated by using the X-SCORE (Wang, Lai, and Wang 2002) (Equation 1). X-score calculates hydrogen bonding, van der Waals, hydrophobic interactions and deformation penalties between the protein and ligand (Obiol-Pardo and Rubio-Martinez 2007).

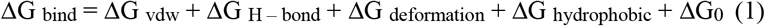

### Molecular Dynamic Simulations

MD simulation were performed by using GROMACS version 2020.4 (Abraham et al. 2015). Gromos53a5 forcefield (Oostenbrink et al. 2004) was used for the simulation. Both protein and protein-ligand complex simulations were performed. Protein was placed in the center of the cubic box with distance of 10A^0^ radius around the protein and SPCE water model was used for the solvation. Ligand topologies were generated by ATB server (Malde et al. 2011). NVT and NPT ensemble were ran for 500 Pico seconds before the minimization step at 300k temperature and 1 bar pressure. 50000 Minimization steps were done before the final production run. Particle Mesh Ewald method (Cerutti et al. 2009) was used for the long-range electrostatic interactions and Lenard Jones potential and coulombs charge was used for the short-range interactions with distance of 1.2 nm. Berendsen thermostat was used for the temperature coupling (Bussi, Donadio, and Parrinello 2008). LINCS algorithm was employed to constrain the bonds with hydrogen atoms. After equilibration, each system setup for the 50ns long production run was performed at 2 femto seconds (fs) time step, for every 10ps trajectory (pdb coordinates are saved) was saved. Results were analyzed by using the inbuilt gromacs modules and results were analyzed and visualized by using XMGRACE tool.

### ADME and Toxicity predictions

Pharmacokinetic (PK) properties namely absorption, distribution, metabolism and excretion (ADME) are the important properties to measure the movement of drug in the body with respect to time. In the present study ADME properties were predicted computationally with the help of ADME descriptor algorithm protocol in BIOVIA Discovery studio 2020 (Accelrys, Inc., San Diego, CA, USA). This algorithm can predict the six properties namely human intestinal absorption, aqueous solubility, Blood Brain Barrier penetration (BBB), plasma protein binding (PPB), CYP2D6 binding and hepatotoxicity of the small molecules. It can also measure the drug likeness properties of the small chemical/biological molecule. Toxicity is one of the important parameters to be check in the animal clinical trials. Once it shows less or no toxicity in the animals, it will go to the human clinical trials. Less toxicity indicates the quality of the chemical substance.

Toxicity of a drug is measure of chemical substance quality by evaluating the damage to the organ in humans/animals and poisonous reactions occurred in the body after drug administration. Toxicity of the drug is predicted computationally by using the Toxicity descriptor algorithm TOPKAT (TOxicity Prediction by Komputer Assisted Technology) in the BIOVIA Discovery studio 2020 (Accelrys, Inc., San Diego, CA, USA). Predicted values of ADME properties are shown in table 5. Ponnan et al. described the levels of ADME properties Intestinal absorption, Aqueous solubility, Blood Brain Barrier penetration, CYP2D6 binding, Plasma Protein Binding and hepatotoxicity according to the values, which are represented below the table 5 (Ponnan et al. 2013). TOPKAT results shows that all the top ten compounds are non-mutagens, when predicted with Ames mutagenesis test.

## Results and Discussion

### High throughput virtual screening of ChEMBL malarial library

As the *P. falciparum* becoming more lethal and resistance is spreading all over the world, there is an urgent need to find the novel molecules with high potency. The selected target PfM17LAP is known to play an important role in the growth of parasite by regulating the release of amino acids. We have done re-docking of reference compound BES present in the protein by defining BES ligand as binding site in LibDock. Screening of ChEMBL antimalarial library was done on the BES ligand binding site. CHEMBL is a manually curated and it is collection of bioactive molecules which are probable drug candidates with drug-like properties(Gaulton et al. 2017). In table 4 we mentioned the bioactivity statistics of top ten LibDock scored compounds. Among the top ten LibDock scored compounds, CHEMBL369831 and CHEMBL191130 had the minimum reported Ki values of 0.4 nM.

Through Insilico screening, we selected top ten LibDock scored compounds as a predicted potential inhibitor, which may inhibit the protein activity by proper binding in the active site. Higher LibDock scores indicate the proper binding of ligand to the target. LibDock score of the reference ligand BES is 130.07. LibDock score of the top scored ligand CHEMBL369831 from the ChEMBL antimalarial database was 230.38. Docked pose of the top scored compound is visualized through Pymol. Figure 1 shows that ligand 369831 in the binding pocket along with the metal ions Zn^+2^ and Mg^+2^ and figure 1B shows Ligplot+ interactions of CHEMBL369831 with protein and metal ions. Compound CHEMBL369831 was making 9 hydrogen bonds and 51 non bonded interactions with protein. CHEMBL176888 was the top 4 ranked LibDock scored compound making 5 hydrogen bonds and 54 non bonded interactions with protein. Detailed scores of the top ten ligands reported in table 2. Post docking analysis with X-SCORE showed that CHEMBL176888 has more binding energy (−11.05 K cal/mol) among the top ten compounds, after that CHEMBL369831 has the highest binding energy -10.9 K cal/mol. X-SCORE binding energies showed that binding affinity of top ten scored compounds were higher than the reference compound BES (−8.75K cal/mol). Binding energies of the top ten compounds along with reference compounds are mentioned in table 2. O45 of the compound CHEMBL369831 was found making hydrogen bonds with LYS374, THR486 and ASP399 as well as coordination with Zn^+2^. O41 of compound CHEMBL176888 was making hydrogen bonds with LYS374, THR486 and coordination with Zn^+2^. O1 of the reference compound BES making hydrogen bonds with ASP399, THR486 and coordination with Zn^+2^. Results showed presence of free terminal oxygen atom that makes more favorable interactions with PfM17LAP. Residue LYS374, ASP399, GLU461, ARG463 THR486, LEU487 and GLY489 were the main residues in the binding pocket. Residues LEU487, THR486, ASP399, LYS386 and LYS374 were the most important interacting residues making hydrogen bonds with top ten ranked compounds.

**Table 2:**
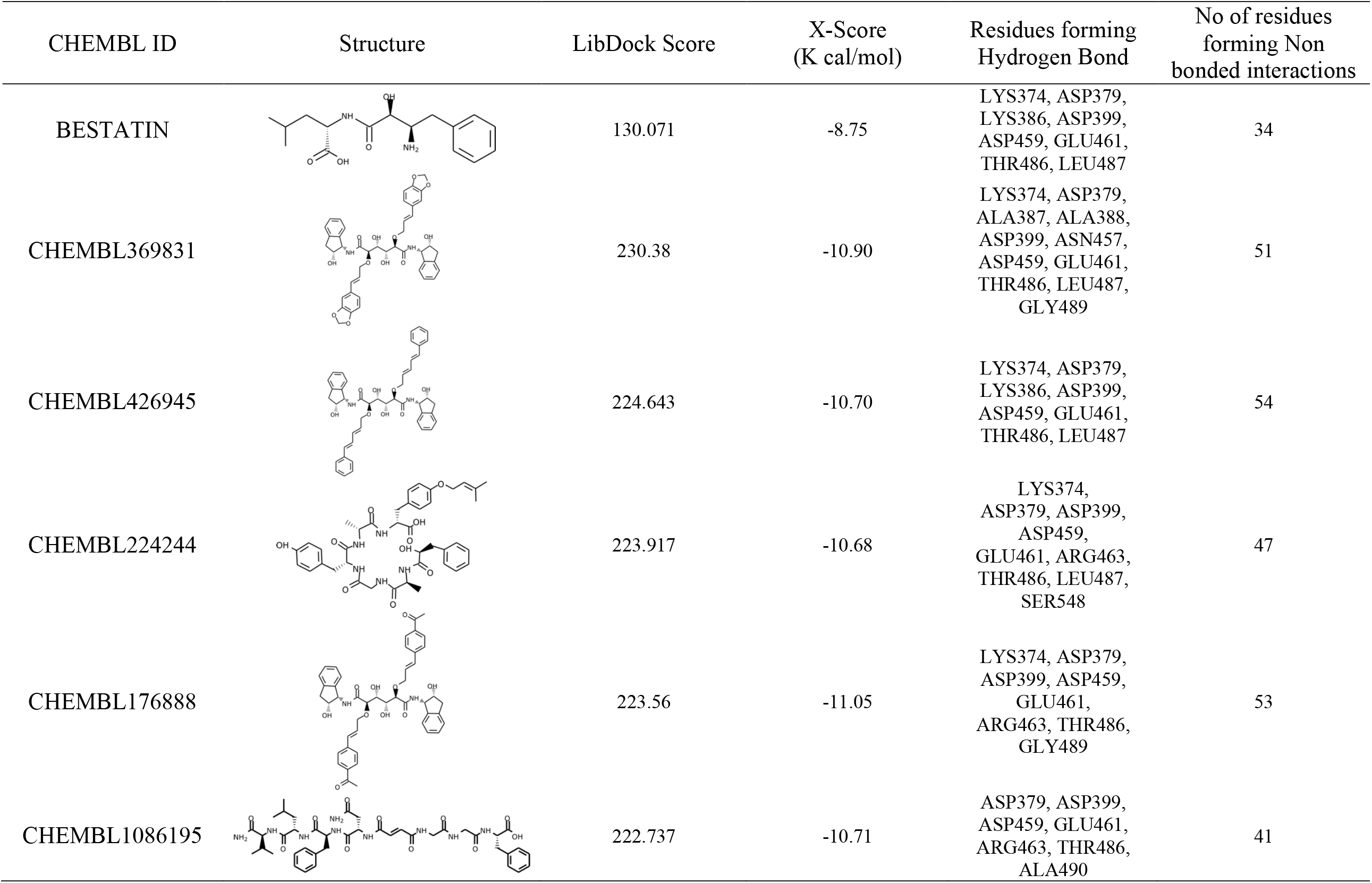
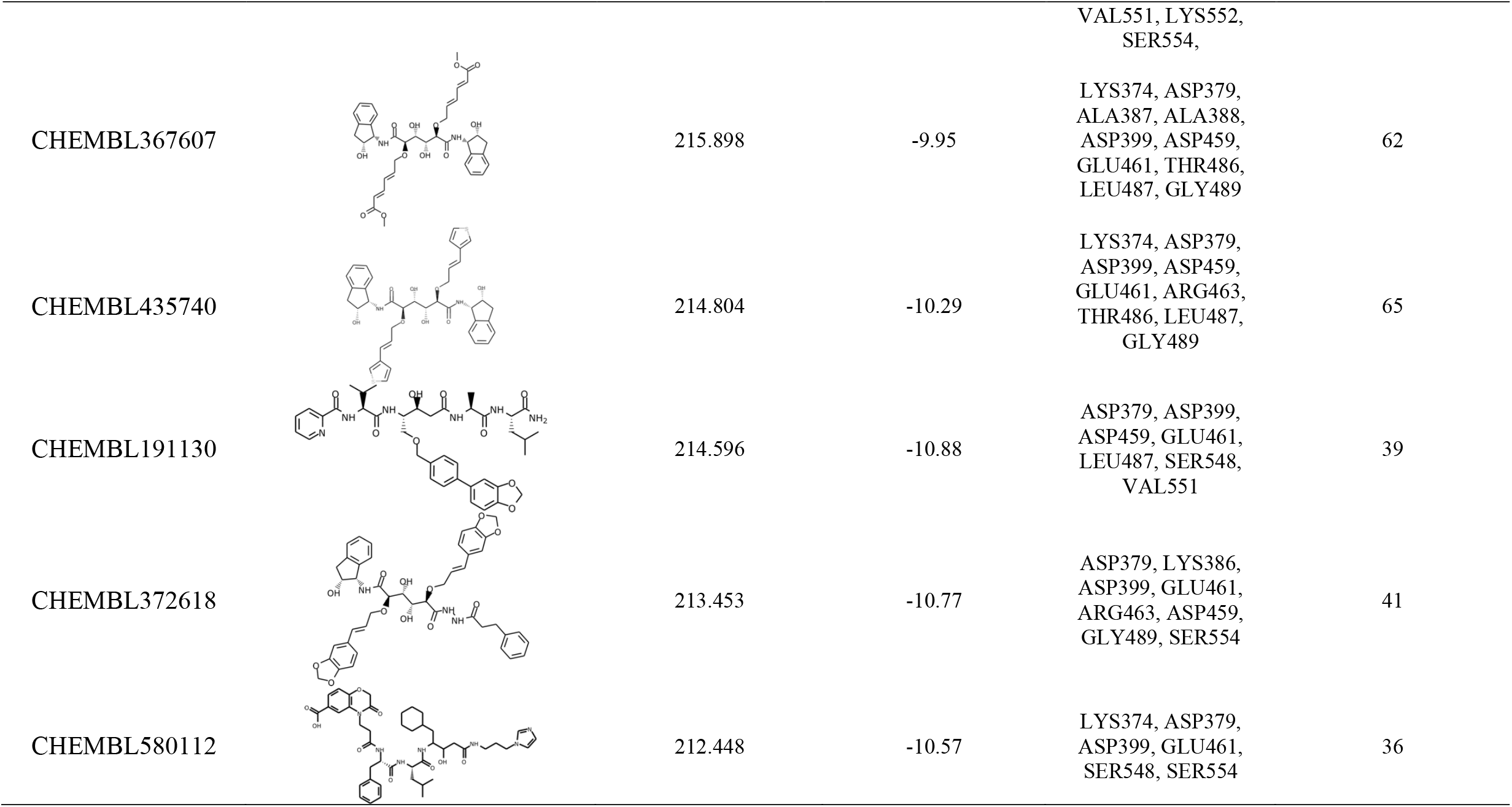
Docking and post docking analysis of the top ten LibDock scored compounds from CHEMBL malarial database, along with reference ligand BES. Post docking analysis done with the help of X-SCORE and Ligplot+ programs.

**Fig 1:**
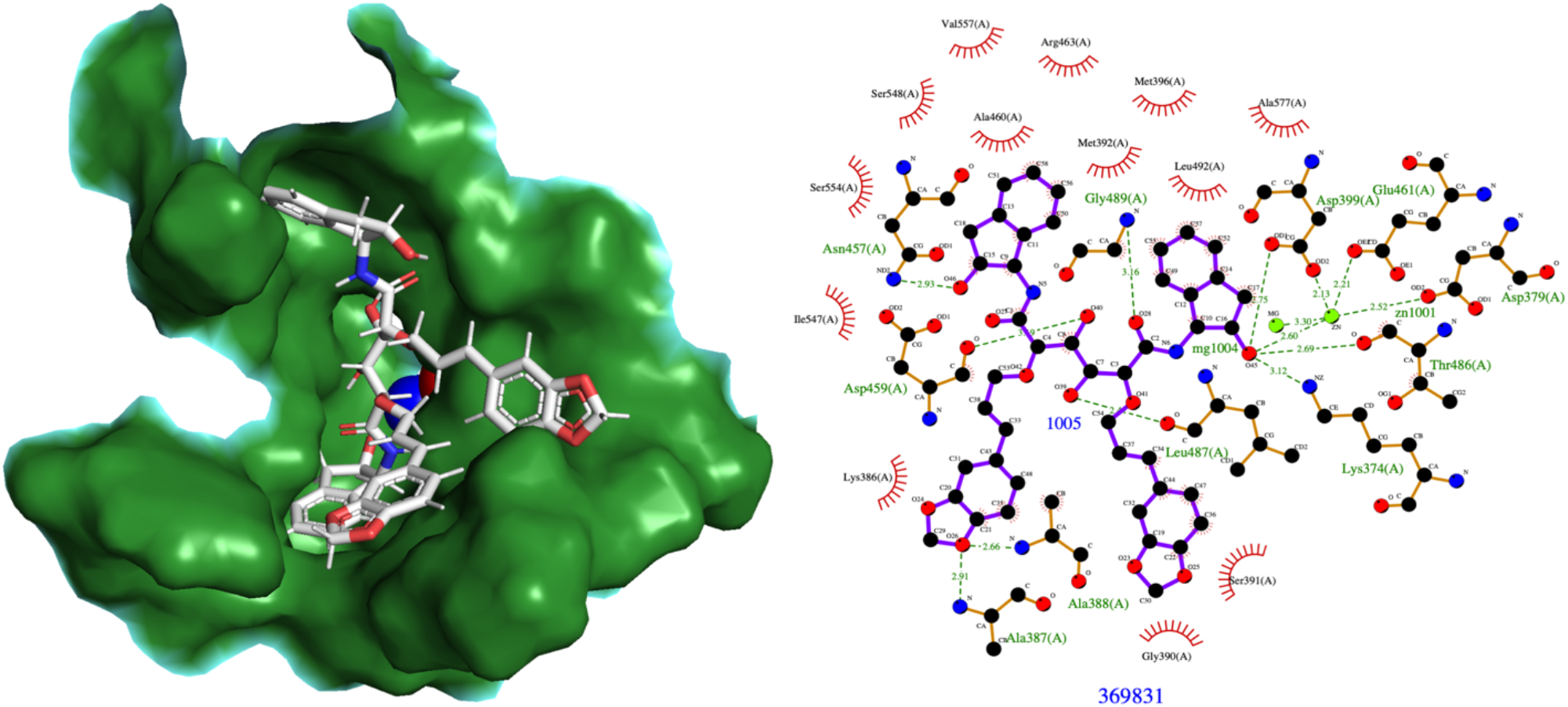
Top scored ligand CHEMBL369831 in the binding pocket of PfM17LAP along with two metal ions Zn^+2^ (Blue) and Mg^+2^ (Red). Ligplot+ interactions of ligand with metals ions and protein.

**Fig 2:**
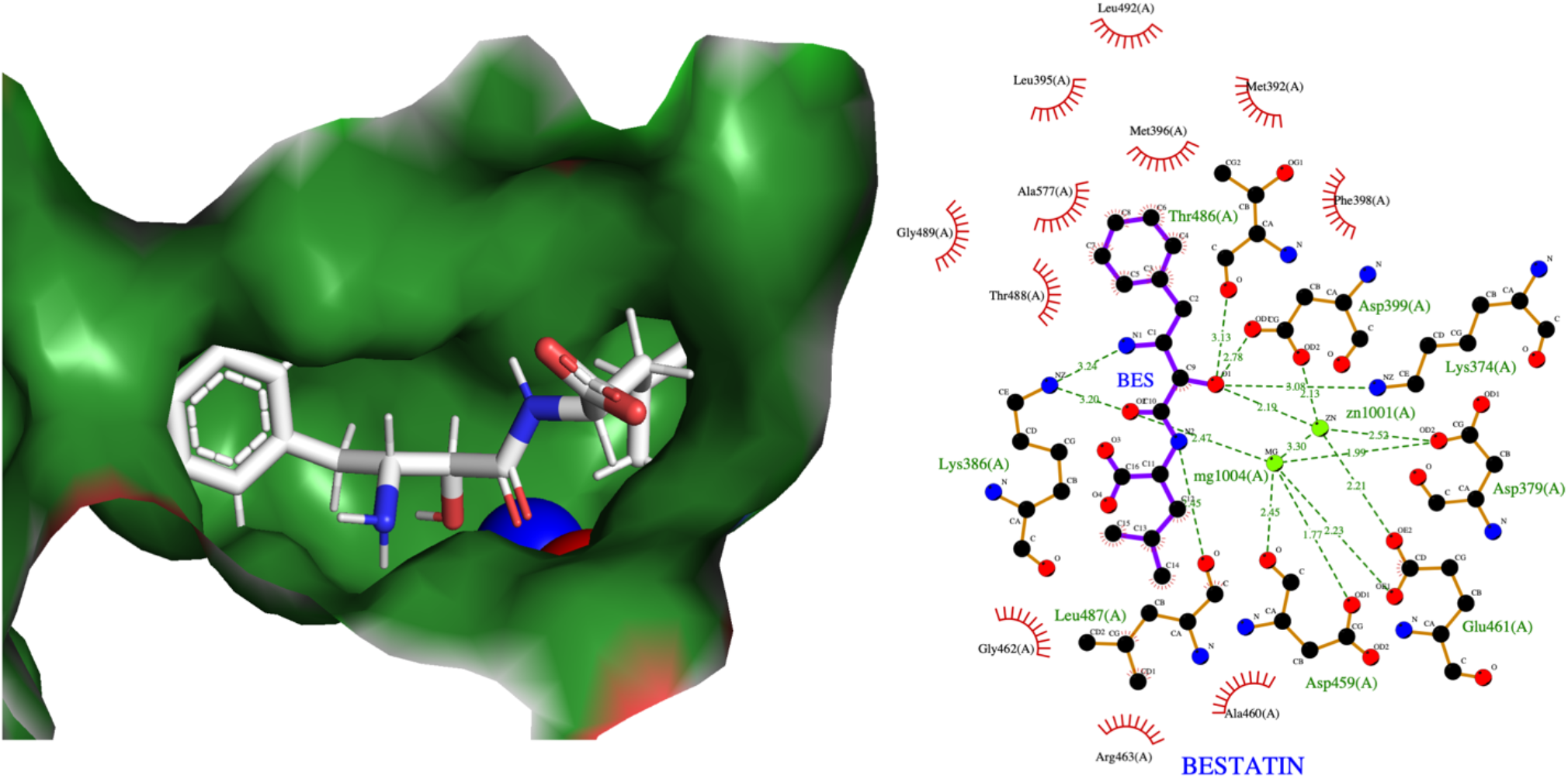
Reference ligand BES in the binding pocket of PfM17LAP along with two metal ions Zn^+2^ (Blue) and Mg^+2^ (red). Ligplot+ interactions of ligand with metal ions and protein.

**Fig 3:**
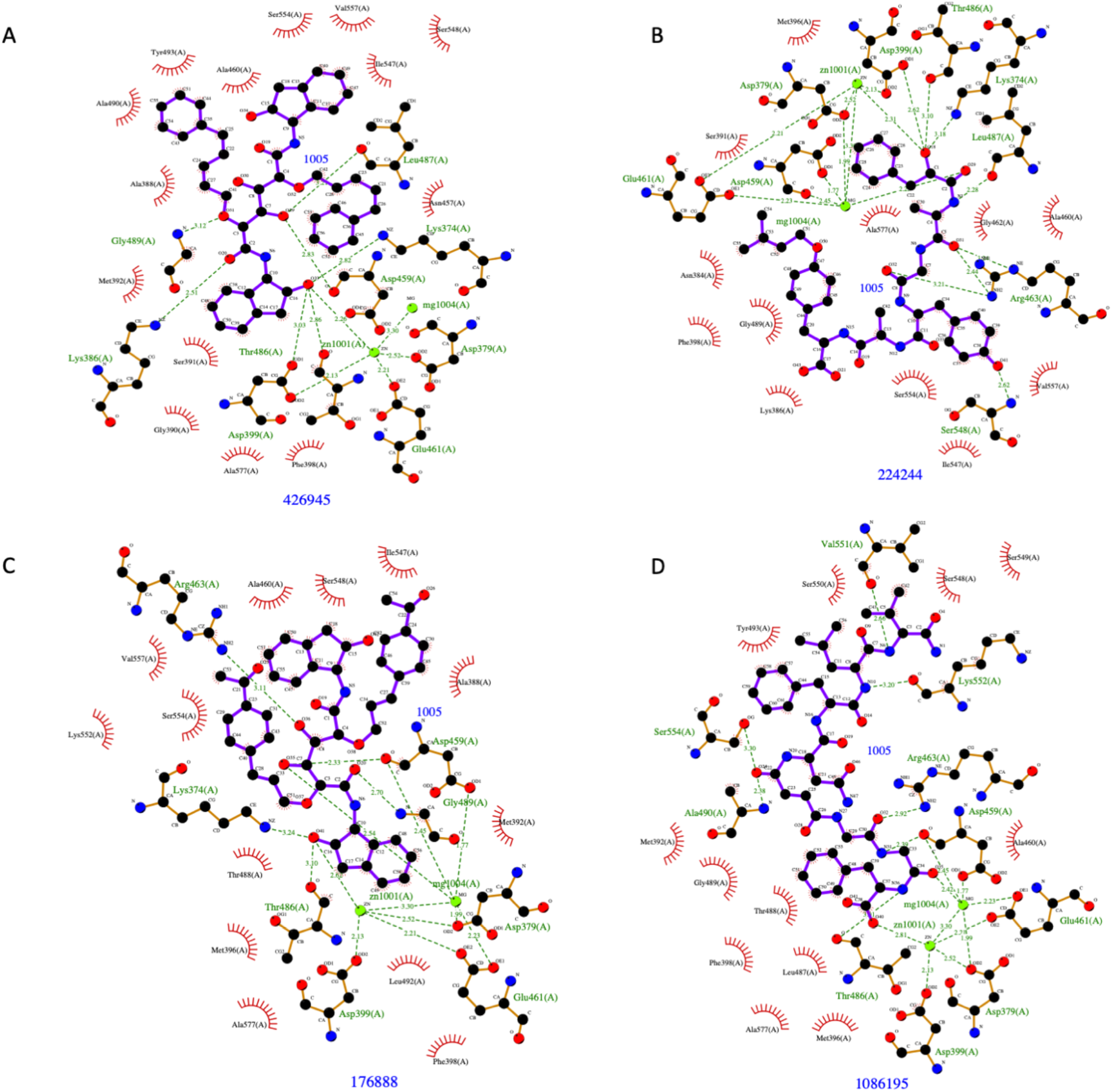
LigPlot+ interaction plots of CHEMBL426945, CHEMBL224244, CHEMBL176888 and CHEMBL1086195 with PfM17LAP. Ligands are represented in violet colour, Zn^+2^ and Mg^+2^ metal ions are represented in green colour. Residues forming hydrogen bonds are labelled in green colour. Bond between two interacting atoms are represented in green colour dotted line along with distance in A^0^.

GLU461, ASP459, ASP399 and ASP379 were most important residues making coordination with metal ions. Overall docking results showed that top scored compounds from ChEMBL library have more affinity than reference ligand BES.

### Molecular docking of Plasmepsins I-IV

Experimental data from the ChEMBL database showed that top ten hits have already been reported to possess inhibitory activity (Ki) against PfPM I, PfPM II and Cathepsin D. Among the top ten hits, 8 compounds have reported high Ki values against the target Plasmepsin I. Therefore, these top hits seem to be acting as probable dual inhibitors. To decipher this, we performed molecular docking analysis of the top ten hits against the four types of plasmepsins found in digestive vacuole. These plasmepsins are known to play an important role in the intraerythrocytic hemoglobin degradation (Nasamu et al. 2020). Docking results showed that the obtained top ten hits maybe effective against the PfPM I-IV. LibDock scores of the top ten hits against PfPM I-IV ranges from 162-213 and X-SCORE ranges from -9.31 to -11.92 Kcal/mol. LibDock and XSCORES of the top ten hits against PfPM I-IV are mentioned in table 3a. LibDock scores of the top ten hits were less for PfPM I-IV as compared to the PfM17LAP. These results suggest that top ten hits maybe more effective towards the PfM17LAP than PfPM I-IV. As the experimental data in table 4 suggest that top hits Ki activity against PfPM-I ranges 0.4 - 4.4 nM and LibDock scores were better for PfM17LAP than PfPM I-IV, theses top hits may show better Ki activity against PfM17LAP. Sequence and structural alignments show no significant similarities between the PfM17LAP and PfPM1 in the active site, still the top ten hits obtained are able to inhibit these two enzymes efficiently along with other plasmepsins. Considering all these results obtained, top ten hits can be considered as potential dual inhibitors.

**Table-3a:** Docking and post docking analysis results of the top ten hits of PfM17LAP against the targets Plasmepsins I-IV. LibDock and XSCORES of the Plasmepsins I-IV are mentioned in the above table.

| S NO | ChEMBL ID | Plasmepsin I LibDock Score | Plasmepsin I X-SCORE K cal/mol | Plasmepsin II LibDock Score | Plasmepsin II X-SCORE K cal/mol | Plasmepsin III LibDock Score | Plasmepsin III X-SCORE K cal/mol | Plasmepsin IV LibDock Score | Plasmepsin IV X-SCORE K cal/mol |
| --- | --- | --- | --- | --- | --- | --- | --- | --- | --- |
| 1 | CHEMBL369831 | 192.962 | -10.74 | 203.242 | -10.52 | 175.422 | -10.62 | 180.552 | -11.11 |
| 2 | CHEMBL426945 | 197.785 | -11.67 | 198.928 | -11.59 | 173.976 | -11.26 | 186.442 | -11.87 |
| 3 | CHEMBL224244 | 171.581 | -9.54 | 182.428 | -10.35 | 168.466 | -9.58 | 170.046 | -9.97 |
| 4 | CHEMBL176888 | 188.765 | -11.77 | 193.115 | -11.92 | 175.588 | -10.24 | 181.078 | -10.97 |
| 5 | CHEMBL1086195 | 190.206 | -10.69 | 168.761 | -10.18 | 170.45 | -9.81 | 200.29 | -10.93 |
| 6 | CHEMBL367607 | 181.611 | -10.05 | 181.826 | -10.51 | 162.381 | -9.78 | 181.714 | -9.31 |
| 7 | CHEMBL435740 | 190.244 | -10.13 | 174.01 | -10.39 | 174.934 | -10.58 | 179.025 | -10.62 |
| 8 | CHEMBL191130 | 168.586 | -10.67 | 176.349 | -11.07 | 168.077 | -10.09 | 179.572 | -10.81 |
| 9 | CHEMBL372618 | 187.873 | -10.59 | 213.478 | -11.02 | 179.378 | -10.44 | 195.931 | -10.97 |
| 10 | CHEMBL580112 | 175.393 | -9.91 | 188.218 | -10.9 | 181.496 | -10.77 | 183.808 | -10.83 |

**Table-3b:** Docking and post docking analysis of the top two hits against the target PfPM-I. Residues making hydrogen bonds with Plasmepsin-I are mentioned in the table. Post docking analysis done with the help of Ligplot+ and X-SCORE.

| CHEMBL ID | LibDock Score | X-SCORE | Residues making hydrogen bonds | No of non-bonded interactions |
| --- | --- | --- | --- | --- |
| CHEMBL369831 | 192.962 | -10.74 | GLY34, TYR189, THR218, ASP290 | 40 |
| CHEMBL426945 | 197.785 | -11.67 | GLY217, ASP32 | 66 |

**Table 4:** Minimum reported Inhibitory activity (Ki) of the top ten hits against the target Plasmepsin I. Bioactivity data taken from CHEMBL database. NA: Inhibitory activity (Ki) data not available for the specified targets

| S.NO | ChEMBL ID | Targets | Inhibitory Activity Ki (nM) (Minimum reported) | Assay Description |
| --- | --- | --- | --- | --- |
| 1 | CHEMBL369831 | Plasmepsin I, Cathepsin D, Plasmepsin II | 0.5 | Inhibitory activity against protease plasmepsin-1 (Plm I) of *Plasmodium falciparum* |
| 2 | CHEMBL426945 | Plasmepsin I, Cathepsin D, Plasmepsin II | 4.4 | Inhibitory activity against protease plasmepsin-1 (Plm I) of *Plasmodium falciparum* |
| 3 | CHEMBL224244 | NA | NA | NA |
| 4 | CHEMBL176888 | Plasmepsin I, Cathepsin D, Plasmepsin II | 0.8 | Inhibitory activity against protease plasmepsin-1 (Plm I) of *Plasmodium falciparum* |
| 5 | CHEMBL1086195 | NA | NA | NA |
| 6 | CHEMBL367607 | Plasmepsin I, Cathepsin D, Plasmepsin II | 3.7 | Inhibitory activity against protease plasmepsin-1 (Plm I) of *Plasmodium falciparum* |
| 7 | CHEMBL435740 | Plasmepsin I, Cathepsin D, Plasmepsin II | 1.3 | Inhibitory activity against protease plasmepsin-1 (Plm I) of *Plasmodium falciparum* |
| 8 | CHEMBL191130 | Plasmepsin I, Cathepsin D, Plasmepsin II | 0.4 | Inhibitory concentration against the plasmepsin-1 of *Plasmodium falciparum* |
| 9 | CHEMBL372618 | Plasmepsin I, Cathepsin D, Plasmepsin II | 6.0 | Inhibition constant against plasmepsin-1 of *Plasmodium falciparum*, 10 min using 3 uM of DABCYL-Glu-Arg-Nle-Phe-Leu-Ser-Phe-Pro-EDANS as substrate |
| 10 | CHEMBL580112 | NA | NA | NA |

**Table 5:** ADME, toxicity and mutagenesis prediction values for reference ligand and top ten scored ligands. Predictions are done with the help of ADME predictor and TOPKAT from Discovery studio. **Absorption**: Human intestinal absorption - 0: good; 1: moderate; 2: poor; 3: very poor. **Solubility**: Aqueous solubility- 0: extremely low; 1: very low, but possible; 2: low; 3: good; 4: very good; 5: extremely good. **BBB**: Blood Brain Barrier penetration- 0: very high penetrant; 1: high; 2: medium; 3: low; 4: undefined. **CYP-2D6**: Cytochrome P450 – 2D6– 0: noninhibitor; 1: inhibitor. **PPB:** Plasma protein binding- 0: weak absorption; 1: high absorption. **Hepatotoxicity:** 0-nontoxic to humans; 1: toxic to humans. **Mutagenicity**: Less negative values indicate no mutagenicity; high negative values indicate more mutagenicity.

| CHEMBL_ID | Absorption | Solubility | BBB | CYP-2D6 | PPB | Hepatotoxicity | TOPKAT Ames Score |
| --- | --- | --- | --- | --- | --- | --- | --- |
| BESTATIN | 0 | 4 | 4 | 0 | 0 | 0 | -12.8 |
| CHEMBL369831 | 3 | 3 | 4 | 0 | 0 | 0 | -19.2 |
| CHEMBL426945 | 3 | 4 | 4 | 0 | 0 | 0 | -13 |
| CHEMBL224244 | 3 | 4 | 4 | 0 | 0 | 0 | -32.2 |
| CHEMBL176888 | 3 | 4 | 4 | 0 | 0 | 0 | -12.2 |
| CHEMBL1086195 | 3 | 5 | 4 | 0 | 0 | 0 | -41.1 |
| CHEMBL367607 | 3 | 4 | 4 | 0 | 0 | 0 | -13 |
| CHEMBL435740 | 3 | 3 | 4 | 0 | 0 | 0 | -8.36 |
| CHEMBL191130 | 3 | 3 | 4 | 0 | 0 | 1 | -23.9 |
| CHEMBL372618 | 3 | 3 | 4 | 0 | 0 | 0 | -23 |
| CHEMBL580112 | 3 | 5 | 4 | 0 | 0 | 0 | -43.1 |

Molecular docking results showed that PfPM I making 4 hydrogen bonds with a distance range of 2.80-3.18 A^0^ and 40 non bonded interactions with the CHEMBL369831. CHEMBL426945 making 2 hydrogen bonds with a distance range of 2.85-2.90 A^0^ and 66 non bonded interactions with PfPM I. Residues making hydrogen bonds with top two hits are mentioned in table 3b and Ligplot+ interactions are shown in figure 4.

**Fig 4:**
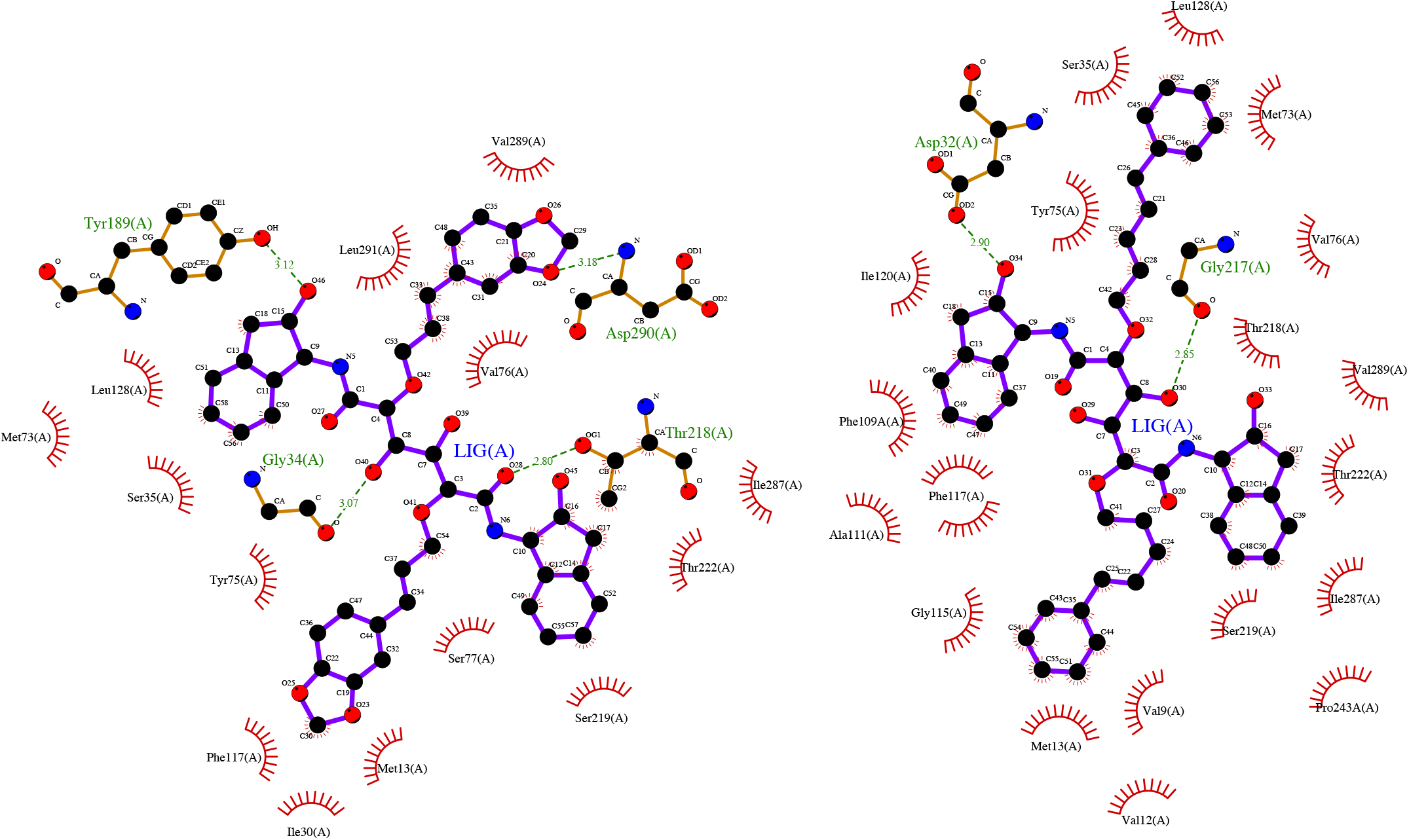
Ligplot+ interactions of CHEMBL369831 and CHEMBL426945 with Plasmepsin I. Residues forming hydrogen bonds are labelled in green colour. Bond between two interacting atoms are represented in green colour dotted line along with distance in A^0^.

**Fig 5:**
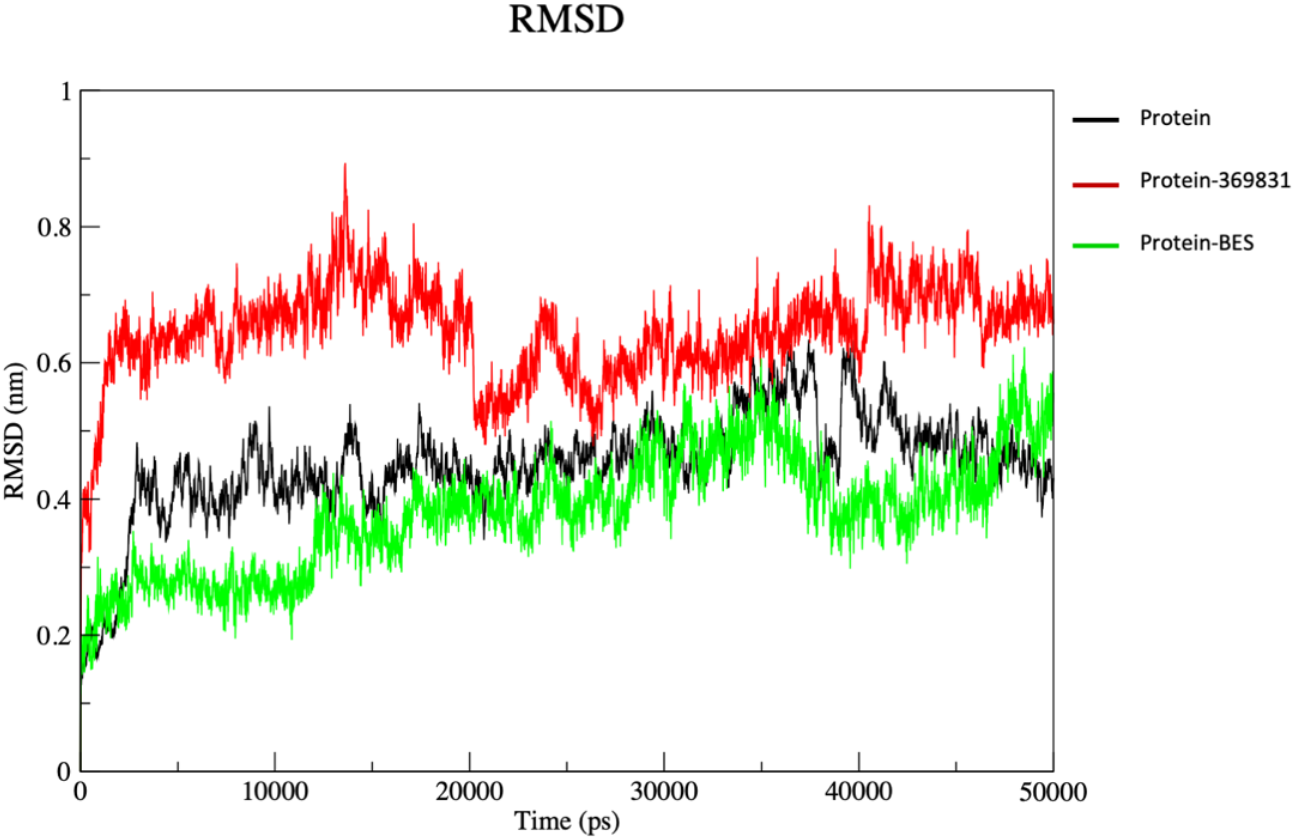
Root Mean Square deviation (RMSD) graph of three systems. Unbound protein, Protein with ligand CHEMBL369831 and protein with ligand BES.

### ADMET and Mutagenesis analysis of top ten scored compounds

ADMET analysis done with the help ADME predictor in Discovery studio. Toxicity predictions are done with of TOPKAT module from Discovery studio. ADME analysis showed that except BES, every ligand shows poor absorption. Solubility level is good for top eight ligands and extremely good for CHEMBL1086195 and CHEMBL5801112. Blood Brain Barrier penetration value is unknown for the top scored ligands including reference ligand BES. As per prediction top ten scored ligands are non-toxic to humans except CHEMBL191130. TOPKAT predictions showed that top scored ligands are non-mutagens.

### Molecular Dynamic simulation analysis

Through MD simulations we can observe the changes in the protein according to time. Root Mean Square Deviation (RMSD), Root Mean Square Fluctuations (RMSF), Radius of Gyration (Rg) and Hydrogen bonds analysis done with the inbuilt modules of the gromacs. Three systems unbound protein, Protein in complex with CHEMBL369831 and Protein in complex with BES are observed for 50ns time. Protein in complex with the CHEMBL369831 shows little deviation in the RMSD in comparison with the other two systems. But the RMSF and Rg of the protein in complex with CHEMBL369831 is stable than the other two complexes (fig6 & fig7). Residues between 328-335 are the part of loop and helix and it is not a part of active site region shows more fluctuations in the protein in complex with BES. Figures 6 shows overall fluctuations of the protein in complex with BES is more as compared to other two systems.

Hydrogen bonds analysis shows that Protein in complex with BES shows a greater number of hydrogen bonds (6) than the other system. Figure 7 shows that protein in complex with the CHEMBL369831 shows 2 consistent hydrogen bonds throughout the simulation.

**Fig 6:**
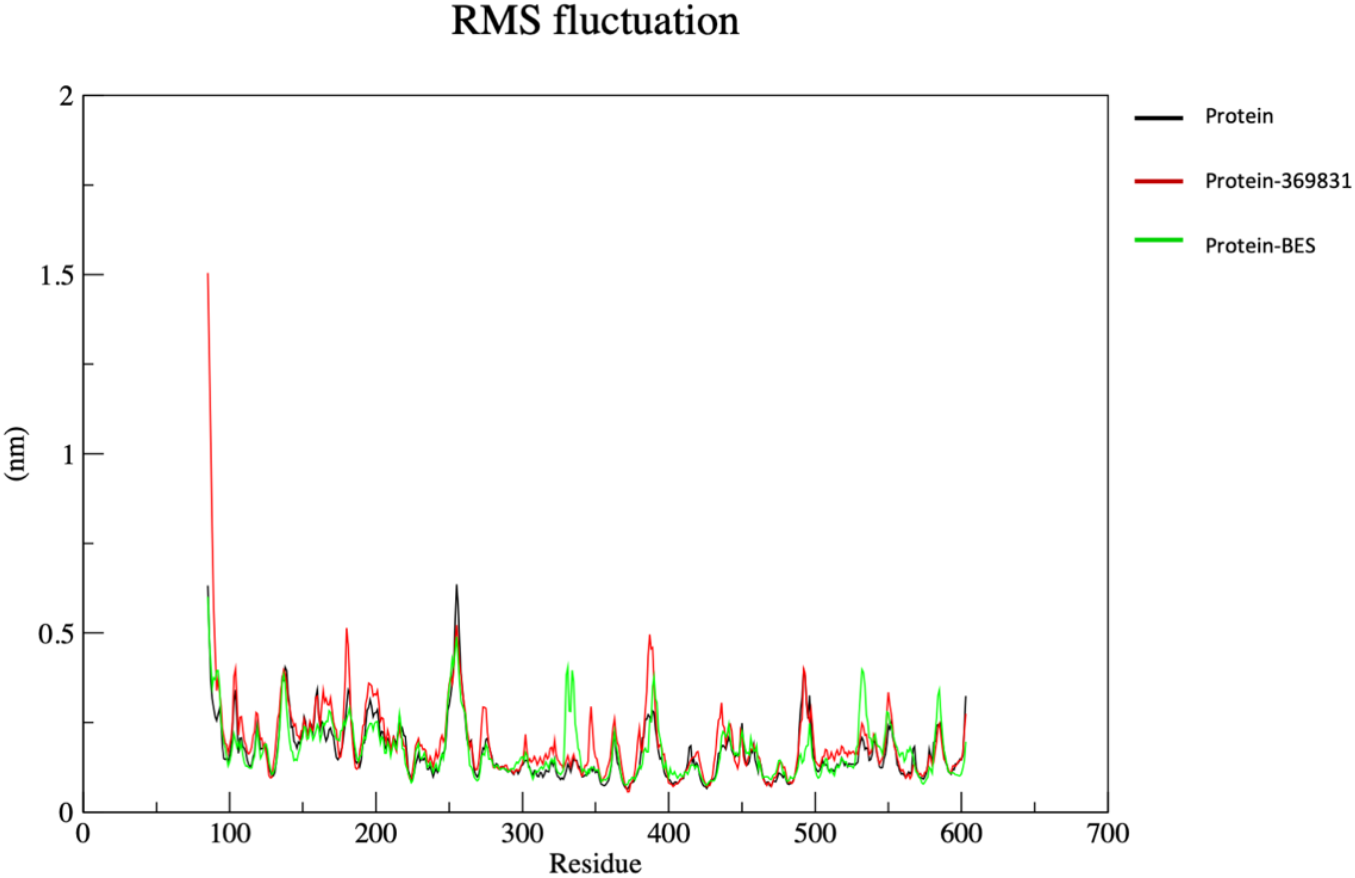
Root Mean Square fluctuation (RMSF) graph of three systems. Unbound protein, Protein with ligand CHEMBL369831 and protein with ligand BES.

**Fig 7:**
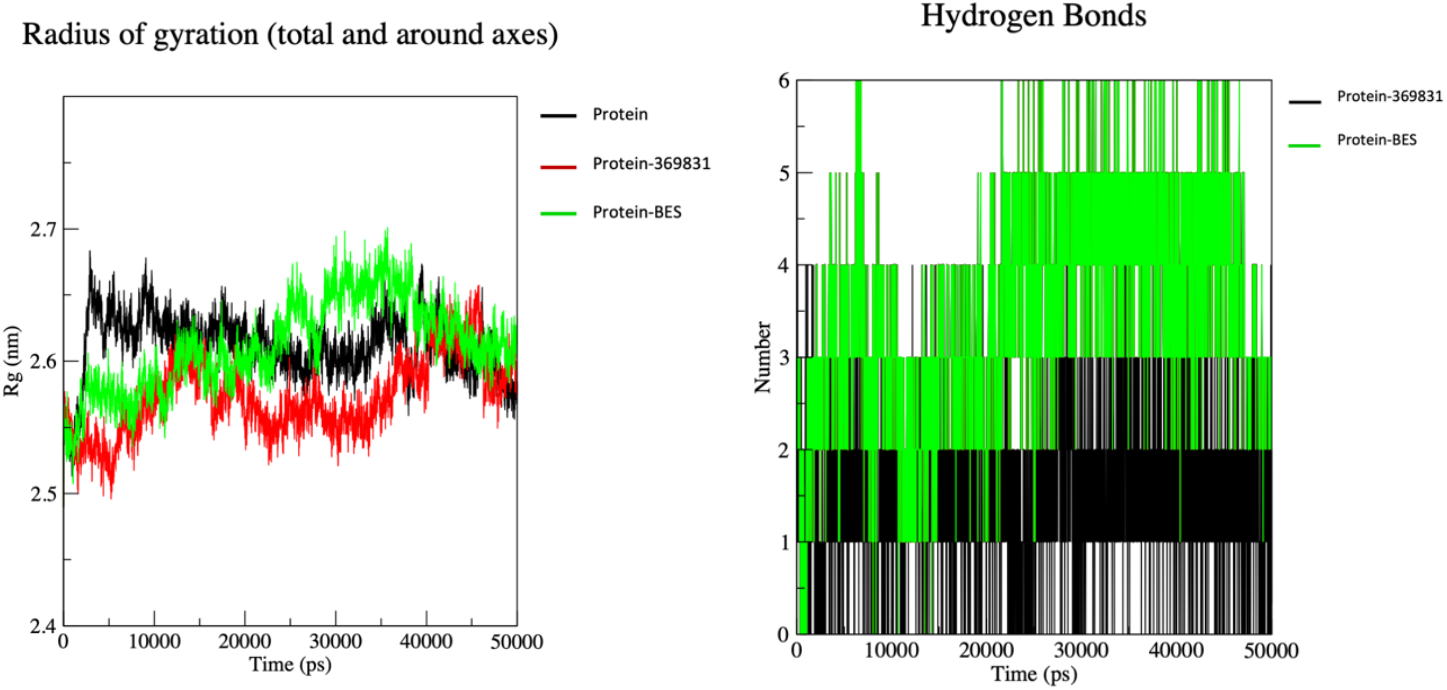
Radius of gyration (Rg) graph of three systems. Unbound protein, Protein with ligand CHEMBL369831 and protein with ligand BES. Hydrogen bonds between protein ligand complexes, Protein-CHEMBL369831 and Protein-BES.

## Conclusion

As resistance to first line antimalarial drugs is increasing rapidly, there is a rising need to find novel antimalarial molecules with grater potency. The aim of our study was to find novel molecules with greater affinity towards the selected target PfM17LAP. It is one of the most crucial enzymes required for the survival of the *P. falciparum*. In the present study, antimalarial database from ChEMBL was taken for the screening purpose whose results showed that CHEMBL369831 and CHEMBL176888 showed greater affinity towards the protein with highest LibDock and X-SCOREs. Further, stability of the protein in complex with CHEMBL39831 was studied by using molecular dynamic simulations. Overall MD simulation results showed that CHEMBL369831 is stable in the active site. Bioactivity data from the CHEMBL database shows that molecule CHEMBL369831 has minimum reported inhibitory constant (Ki) of 0.5 nM. Overall results showed that CHEMBL369831 may be the potential inhibitor for PfM17LAP. Further, clinical studies are required to use this molecule as an antimalarial drug. The experimental data from ChEMBL database and inferences drawn from our present investigation, together suggest that the top ten hits seem to act as dual inhibitors of Plasmepsins I-IV and PfM17LAP.

